# Mapping of snail intermediate host habitat reveals variability in schistosome and nonschistosome trematode transmission in endemic settings

**DOI:** 10.1101/2023.06.04.543635

**Authors:** Teckla Angelo, Naima Camilla Starkloff, Moses Paul Mahalila, Jenitha Charles, David James Civitello, Safari Kinung’hi

## Abstract

**Background:** There is growing recognition that mass drug administration must be complemented with environmental interventions to interrupt schistosomiasis transmission. Accurate mapping of snails and schistosome parasite distribution is critical to identify foci of human exposure and prioritize sites for interventions.

**Methodology:** We conducted longitudinal environmental surveys of snails and schistosomes in 467 waterbodies across 86 villages in northwestern Tanzania to describe spatial and temporal variation in snail and schistosome parasites presence and identify relevant underlying predictors. We conducted time constrained net sampling of *Bulinus* snails from vegetation, sediments, and floating objects and then examined all collected snails for patent infections.

**Principal findings:** A total of 43,272 *Bulinus* snails were collected across the three visits to each waterbody spanning November 2020 – August 2021, and we conducted statistical analyses on the latter two visits with more in-depth surveys (25,052 snails). We found patent schistosome infections in 0.87% of snails, 9.8% of waterbodies, and 31% of villages in all six districts. Variance decomposition indicated that variation among waterbodies was associated with variation in snail presence and the prevalence of schistosomes and nonschistosome parasites, whereas variation among villages and districts was not. Snail presence was highest in March-May a period of heavy rains, but otherwise not associated with waterbody characteristics. Waterbodies permitting cattle use had significantly higher prevalence of schistosomes than those permitting only human use. Nonschistosome parasites were more prevalent in June to September the dry season but were not associated with other waterbody characteristics. Waterbody permanence and distance to the nearest primary school were not associated with snails or parasites.

**Conclusions/significance:** This study revealed substantial variation in snails, schistosome and nonschistosome abundance at local (waterbody) scales, and it suggests links between community-driven water use decisions and schistosome transmission. The identification of local drivers of snail and schistosome abundance level and transmission factors at waterbody scale can complement studies across larger scales to shed light on transmission hotspots and guide the development of targeted interventions for schistosomiasis control.

**Author summary:** Currently there is increasing need to supplement mass drug administration with environmental interventions by identifying potential sites for human exposure to disrupt schistosomiasis transmission. Here we focus on environmental surveys to determine leading factors for presence and sequential variation in snails and schistosomes. Using a timed sampling strategy, snail vectors were collected and examined for infections. We screened snails collected from November 2020 to August 2021 and found variation among waterbodies in snail presence and infections of schistosomes and nonschistosome parasites. This variation was not observed at village and district levels. High abundance of snails was observed in March to May during heavy rainfall but not connected with waterbody distinctiveness. Waterbodies used by cattle had higher prevalence of schistosomes compared to waterbodies used by humans alone. Nonschistosomes were higher in June to September the dry season but not related to waterbody characteristics. Although chemotherapy reduces schistosomiasis burden, our results suggest that identification of transmission sites at waterbody scale could aid development of targeted interventions for schistosomiasis control.

## Introduction

Schistosomiasis is a tropical disease of medical and veterinary importance caused by trematodes in the *Schistosoma* genus that cycle between snail intermediate hosts and mammal definitive hosts, including humans (1,2). It is a disease of the poor and ranks second next to Malaria in terms of disease burden in Sub-Saharan Africa (3,4). More than 200 million people are infected globally, while 440 million are at risk of infection(5). Global efforts to control schistosomiasis using mass drug administration (MDA) of the anthelminthic praziquantel have substantially reduced schistosomiasis morbidity in humans (6,7). However, there is growing recognition that environmental interventions are required to disrupt transmission and facilitate schistosome eradication(8).

Environmental interventions, such as chemical or biological control of intermediate host snails (6,9,10), depend on parasitological mapping of snails and schistosomes within and across schistosome-endemic and at-risk communities. Human parasitological surveys of schistosome infection often span entire communities or cohorts of school-aged children. Humans contract schistosomiasis in freshwater sites, and exposure can be extremely heterogeneous even within a village(11,12). Members of villages may be likely to convene on certain sites, such as around primary schools, yet also disperse to other sites more heterogeneously, e.g., those nearest to individuals’ homes for domestic water collection. Thus, accurate mapping of snails and schistosomes can identify foci of human exposure, prioritize sites for interventions, and connect with human parasitological data to determine how environmental interventions translate into human health gains.

Schistosome and snail abundance can vary dramatically across spatial and temporal scales. At the national and regional scale, human infection with *Schistosoma mansoni* is greatest near large, permanent waterbodies, such as Lake Victoria, whereas *S. haematobium* prevalence is greatest in communities accessing smaller, often seasonal waterbodies(13). This broad-scale pattern reflects contrasting habitat use by *Biomphalaria* snails, which predominate permanent sites and transmit *S. mansoni* a causative agent of intestinal schistosomiasis, and *Bulinus* snails, which predominate temporary sites and transmit *S. haematobium* a causative agent of urogenital schistosomiasis in humans(14,15).Across villages, schistosome infection prevalence also varies widely, reflecting differences in MDA history, ecological conditions, epidemiological factors, and social or behavioral norms (16,17). At the scale of individual waterbodies, snail populations can vary seasonally, with waterbody ephemerality and other characteristics, such as waterbody size, coverage of floating vegetation, physical and chemical water parameters, and whether the site is manmade(18,19). Additionally, landscape features of water bodies, such as close proximity to households and schools is more likely to harbor infected snail intermediate hosts which are transmission sources of schistosomiasis to inhabitants(20,21).Achieving significant reduction of schistosomiasis burden by identifying potential hotspots of transmission would complement the ongoing human treatment and attain schistosomiasis elimination goals based on the WHO road map(22).

We conducted repeated environmental surveys of snails and schistosomes in 467 waterbodies across 86 villages in Northwestern Tanzania to characterize spatial and temporal variability in snails and schistosomes and identify predictive habitat characteristics. Villages in our study area are inland from Lake Victoria, and human schistosome transmission occurs as humans interact with relatively small public water bodies in their villages, due to limited access to running water(5,23). Additionally, these public water sources are used by both humans and animals, predisposing them to schistosome infections(20). These water sources harbor snail intermediate hosts that transmit both human and animal parasites, specifically *Bulinus spp.* snails that can transmit both *Schistosoma haematobium* (infecting humans) and *S.bovis* (infecting cattle). These two trematode species have been documented to hybridize in sympatric transmission settings(24,25). *Bulinus* species also harbor other cattle trematode parasites (non-schistosome) such as different types of Xiphidiocercaria and *Strigea* (26). In this study, we specifically 1) assessed snail presence as well as human schistosome and cattle trematode infections in water bodies located in six Tanzanian districts in the Lake Victoria, 2) examined the variability in infection prevalence at three spatial scales (waterbody, village, and district), and 3) evaluated the associations between waterbody characteristics such as size, restriction of cattle use, school proximity and season on schistosome and non-schistosome prevalence. Understanding the factors that promote the transmission of both human and livestock parasites is crucial for more precise and sustainable control and elimination of schistosomiasis.

## Methods

### Sampling sites

This study served as a geographically broad and locally intensive mapping program of snail presence and infection. Six districts (Magu, Misungwi, Kishapu, Busega, Sengerema and Kwimba) in Mwanza, Simiyu and Shinyanga regions of Northerwestern Tanzania were included in this study (Fig 1A). Within these six districts, 86 villages were selected in which 467 water bodies (sampling sites) were surveyed for *Bulinus* snails from November 2020 to August 2021. Community leaders participated in snail sampling at each water contact site in their respective villages. The sampling sites were located further inland from Lake Victoria and ranged from small, temporary to large, permanent water bodies. Each of the sampling sites was surveyed one to three times within three different sampling phases during the transmission season, Phase 1 (November 2020 to February 2021; short rainy season), Phase 2(March to May 2021; heavy rain season), and Phase 3(June to August 2021; dry season) (Fig 1B). The GPS coordinates of each water body was recorded each visit.

**Fig 1.**
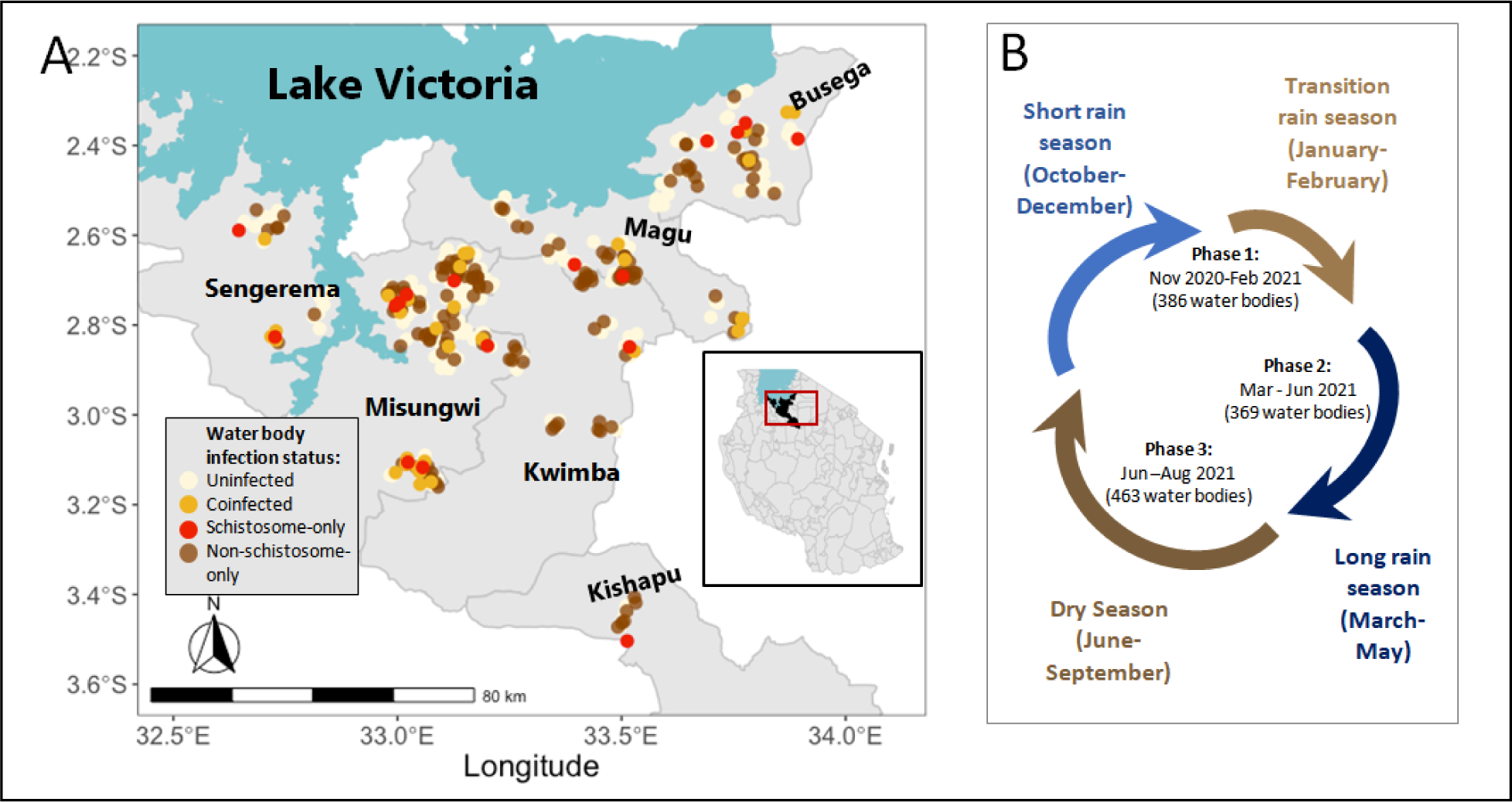
Map of the study area showing snail infections and seasonal patterns in Northwestern Tanzania (A) Map of waterbodies surveyed in the Lake Victoria watershed for schistosome (phases 1-3) and cattle-trematode (phases 2-3) infected *Bulinus* snails, with colors indicating the infection status of the waterbody (uninfected, singly infected by either parasite species or coinfected by both parasite species). Inset shows position of the six districts sampled in the Lake Victoria watershed of Northwestern Tanzania. (B) The local rainfall seasonality in Tanzania (27) and the timing of snail sampling within these seasons.

### Snail sampling

The variability in snail presence and infection prevalence of schistosomes and non-schistosomes parasites in snail vectors was assessed in each waterbody inland from Lake Victoria. Three species of *Bulinus (B. nasutus, B. africanus* and *B. globosus)* were of interest as potential intermediate hosts for *S. haematobium,* the causative agent of urogenital schistosomiasis in humans and S*. bovis,* the causative agent of bovine schistosomiasis in cattle. Other snail species namely, *Pila, Melanoides, Biomphalaria, Lymnaea, Cleopatra, Ceratopharus* and *forsikalii* available at each waterbody were recorded and a small number were saved as species representatives. Snails were sampled using a standardized time-constrained net sampling procedure(28) whereby two researchers collected snails using metal mesh scoop nets, from vegetation and objects floating in water for 15 minutes per site. Collected snails were placed in clean plastic containers and transported to the laboratory of the National Institute for Medical Research (NIMR), Mwanza-Center for examination of patent cercarial release.

### Snail habitats and correlation with snail infectivity

At every waterbody, the presence of vegetation cover was recorded as well as season of collection, substrate and presence of livestock or indication of livestock visits to the sampling sites. Other factors such as water contact by humans and domestic animals, type of water body (being pond, stream, river, dam or natural well), nature of water body (permanent or temporary) and the size of the water body were recorded.

### Assessment for trematode infections

In the NIMR Mwanza Centre laboratory, *Bulinus* snails collected were cleaned and counted. Patent infections were determined using a 24-hour shedding method(29). Snails were placed in 30ml beakers for 24 hours in natural light conditions and, thereafter, were assessed under a dissecting microscope for the presence of larval trematode cercariae.Emerging trematodes with a fork-tailed morphology indicated schistosomes. This morphological diagnostic method lumps together *S. haematobium, S. bovis,* and any schistosome hybrids emerging parasites without forked-tails were classified as non-schistosomes, likely cattle or bird infecting parasites (30).

### Statistical Analysis

All snails collected in phases 2 and 3 were analyzed for infection with schistosomes and other trematode parasites using the generalized linear mixed effects models (GLMMs). Snails from phase 1 were excluded due to lack of records for nonschistosome parasite infections. In all cases, the generalized linear mixed model (GLMMs) with binomial error distributions was used. Before considering any predictors, variance and estimated proportion of total variance was partitioned in each endpoint and explained across spatial scales (district, village, and waterbody) by fitting a random effects-only model with random effects of district, village, and waterbody using the rpt function in the rptR package (31). Because the vast majority of variance occurred at the waterbody scale, the explanatory ability of several waterbody-specific fixed effects: permanence (permanent vs. temporary), longest dimension (in meters), distance to school (in kilometers), permission for cattle use (human use only vs. cattle use permitted), and phase (Phase 2 or 3, see Fig 1B) was tested. While data was collected on schistosome infections across all three seasons, data on non-schistosome infections were recorded in Phase 2-3. The longest dimension data was also limited to the latter two phases. As a result, all statistical models only included data from these two later phases. The GLMMs was fit using the glmmTMB function in the glmmTMB package(32,33). In all statistical analyses, water bodies which were completely dry were excluded. For all snail infection analyses, water bodies in which no snails were found were also excluded.

### Ethical statement

The study received ethical approval from the Medical Research Coordination Committee (MRCC) of the Tanzania National Institute for Medical Research (NIMR), ethics approval certificate number NIMR/HQ/R.8a/Vol.IX/3462.

## Results

### General patterns of snail presence and infectivity

In this study, a total of 43,348 *Bulinus* snails were collected across the three phases (Fig 1). However, only 25,052 snails that were shed for both trematode types between March – August 2021 (Phase 2 and 3) were statistically analyzed from 388 waterbodies in six districts. Snails were abundant in all districts, with the highest number collected in Magu district. Infection rates of nonschistosome parasites were consistently higher than those of schistosome parasites, regardless of spatial scale. Overall, 0.87% of *Bulinus* snails were patently infected with schistosomes, whereas 4.9% were shedding nonschistosome parasites. Co-infections were incredibly rare, with only 0.05% of snails being infected by both groups of trematodes. Similar rates were seen at the village and district level (Table 1).

**Table 1:** Patterns of snail presence and infectivity by waterbody, village, and district

|  | District |  |  |  |  |  | 217 |
| --- | --- | --- | --- | --- | --- | --- | --- |
|  | Busega | Kishapu | Kwimba | Magu | Misungwi | Sengerema | TOTAL |
| Villages | 17 | 3 | 4 | 13 | 27 | 7 | 71 |
| Waterbodies | 62 | 10 | 26 | 87 | 175 | 28 | 388 |
| Schools | 27 | 3 | 4 | 18 | 31 | 9 | 92 |
| Schistosome infected waterbodies (%) | 7<br>(11.3%) | 1<br>(10%) | 2<br>(7.7%) | 5<br>(5.7%) | 18<br>(10.3%) | 5<br>(17.9%) | 38<br>(9.8%) |
| Non-schistosome infected waterbodies (%) | 26<br>(41.9%) | 7<br>(70.0%) | 11<br>(42.3%) | 39<br>(44.8%) | 86<br>(49.1%) | 11<br>(39.2%) | 180<br>(46.4%) |
| Co-infected waterbodies (%) | 1(0.02%) | 0 (0%) | 1 (0.04%) | 0 (0%) | 6(0.03%) | (0.07%) | 10(3.0%) |
| Bulinus snails | 4081 | 786 | 1430 | 6305 | 11283 | 1167 | 25052 |
| Schistosome infected snails (%) | 89<br>(2.2%) | 2<br>(0.25%) | 3<br>(0.21%) | 8<br>(0.13%) | 102<br>(0.90%) | 15<br>(1.3%) | 219<br>(0.87%) |
| Non-schistosome infected snails (%) | 177<br>(4.3%) | 32<br>(4.0%) | 53<br>(3.7%) | 374<br>(5.9%) | 549<br>(4.9%) | 43<br>(1.9%) | 1228<br>(4.9%) |
| Co-infected snails (%) | 1<br>(0.02%) | 0<br>(0.0%) | 1<br>(0.07%) | 0<br>(0.0%) | 8<br>(0.07%) | 2<br>(0.17%) | 12<br>(0.05%) |

The number of villages within which waterbodies were sampled, the number of waterbodies sampled and number of schools georeferenced in each district. For each district, the numbers of waterbodies and snails singly infected by each parasite species or those co-infected with both parasite species were also quantified. These data were collected in phases 2 and 3 and used for all analytical models.

Snail presence and infection status with both schistosomes and non-schistosome parasites showed the greatest variation at the waterbody scale, but not at the larger spatial scales. Specifically, 17% of the total variance in snail presence/absence was associated with waterbodies, 1.2% was associated with villages, and 0% associated with district (Table 2). Schistosome infection prevalence showed the strongest associations with waterbody-scale variation (57% of total variance vs. ∼9.5% for non-schistosome parasites), and infection prevalence for both groups was essentially unassociated with village- and district level variation (∼0% variance at both levels for both parasite groups) (Table 2).

**Table 2:** Proportion of total variance explained in at three spatial scales in random-effects only models.

| Proportion of total variance associated with each spatial scale: |  |  |  |
| --- | --- | --- | --- |
| Endpoint | Waterbody | Village | District |
| Snail presence | 0.17 | 0.012 | 0.004 |
| Schistosome infection prevalence | 0.57 | 0.0035 | 0 |
| Non schistosome infection prevalence | 0.095 | 0 | 0 |

### Factors associated with snail infection at the waterbody scale

Schistosome infection occurred at significantly higher prevalence in waterbodies with permission for cattle use than those for human-use only (p=0.00859, Fig 2a). However, nonschistosome trematode parasite prevalence did not vary with permission of animal waterbody use (p>0.05, Fig 2b). Nevertheless, the prevalence of non-schistosome parasites was significantly higher in Phase 3 than Phase 2 (p<0.0001, Fig 3a). The probability of snail presence was significantly higher in Phase 2 than in Phase 3 (P<0.001, (Fig 3b). No other factors were significantly associated with snail presence. Neither parasite group (Schistosome or nonschistosome) varied in prevalence with waterbody permanence or distance to closest school (p>0.05).

**Fig 2.**
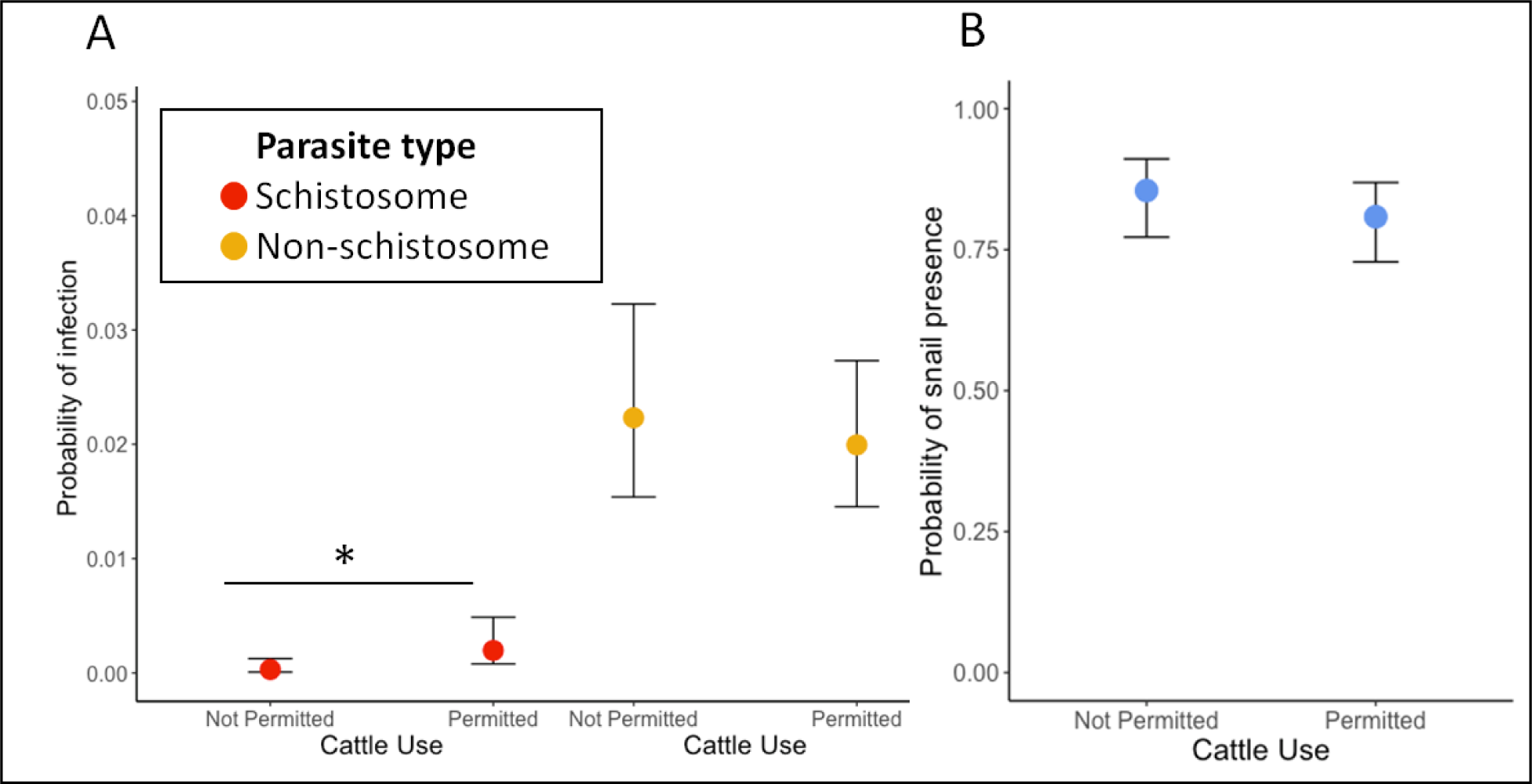
Infection and snail presence with cattle use permission (A)The variability in the probability of infection in waterbodies with and without cattle use permission of schistosome (red) and nonschistosome (yellow) parasites. Schistosome infection is significantly higher in waterbodies where cattle use is permitted than not permitted (p=0.00859), whereas there is no difference with nonschistosome parasites infection (>0.05). (B) Probability of snail presence (blue) does not vary with cattle use permission (>0.05).

**Fig 3.**
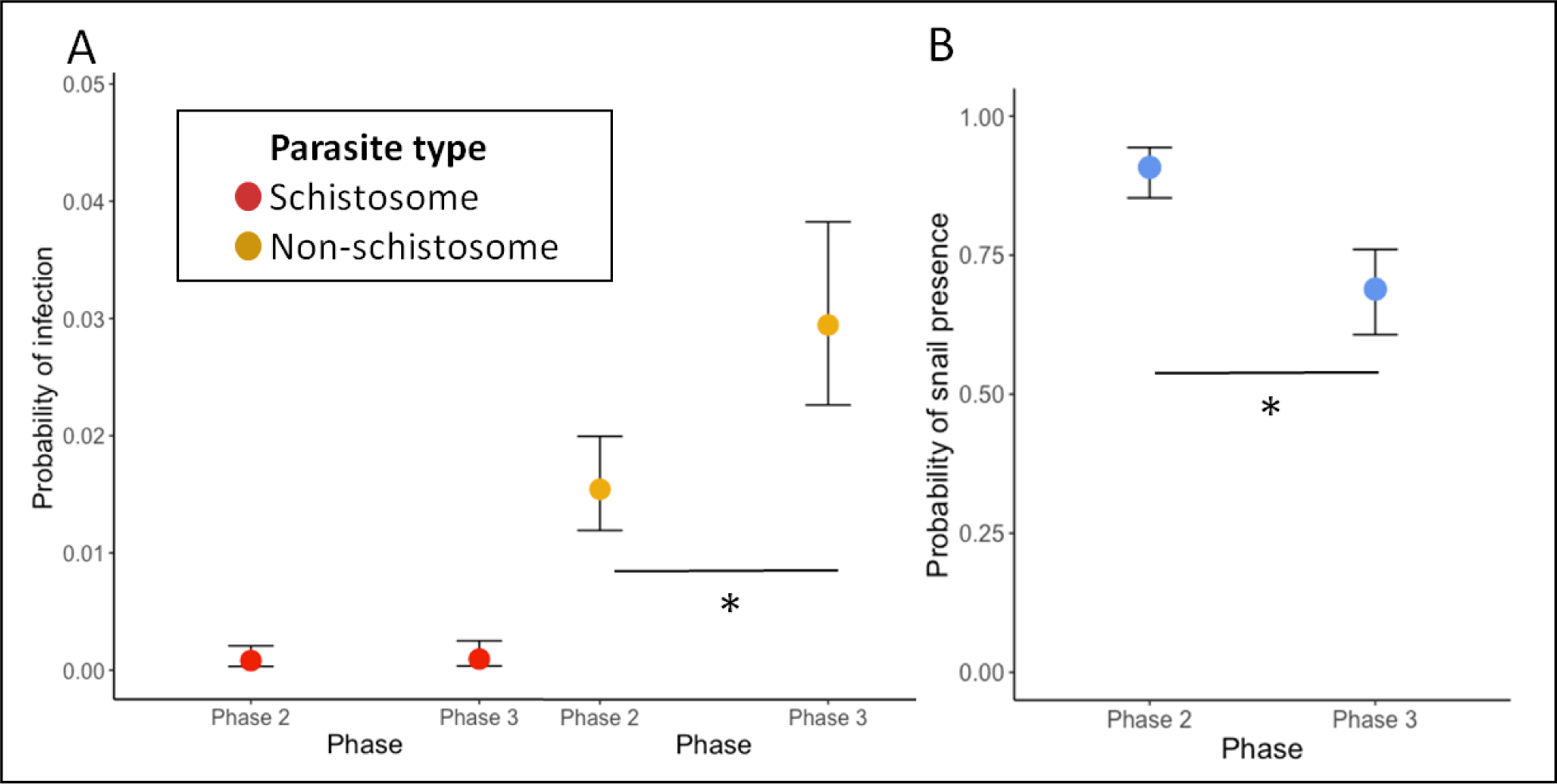
Infection and snail presence with phase of study (A) Variability in probability of infection in waterbodies across the two sampling phases of schistosome (red) and nonschistosome (yellow) parasites. Nonschistosome infection is significantly higher in phase 3 (p<0.001), whereas there is no difference with schistosome infection (>0.05) and increased significantly with waterbody maximum length (p=0.000328 presence (blue) is higher in phase 2 than phase 3 (>0.001). (B) The probability of snail presence (blue) is higher in phase 2 than phase 3 (>0.001).

## Discussion

We investigated the presence of snails, schistosome and nonschistosome trematode parasites across a heterogeneous landscape in Northwestern Tanzania. In our survey spanning six districts, we found that snails, schistosomes, and other trematode parasites are highly variable across space. However, this variability was particularly concentrated at the waterbody scale. This concentration of variation at the waterbody scale suggests that specific activities, preferences, or water uses determine which waterbodies become infected by schistosomes or nonschistosomes parasites. For example, the schistosome association with cattle use permission could indicate a direct role for cattle as well as the presence of *S. bovis* or hybrids, but it could also be associated with human behavior changes in such waterbodies. Similarly, the association between waterbody size and nonschistosome parasites could indicate a direct effect of waterbody permanence or indirect mediation of social factors or behaviors that depend on waterbody size.

While snails were more likely to be present later in the season (Fig 3B), we did not identify any other factors explaining snail occurrence. Other studies have demonstrated that schistosome-vectoring snails can be associated with water parameters, such as ionic concentration (34), water permanence and flow rate (18), aquatic vegetation (19),presence or absence of predators(10). A deeper investigation of aquatic vegetation and predators among these waterbodies is therefore recommended.

The high variability of schistosome prevalence at the waterbody scale has implications for prevention of exposure and transmission control. First, it suggests that most villages contain some hotspots and coldspots of transmission potential, rather than relatively uniform transmission potential across waterbodies. While we documented this variation within a year, multi-year studies are critical to determine if hotspots of transmission persist through time or pop up sporadically. If hotspots are relatively stable, then interventions and/or avoidance behaviors could be directed at a subset of waterbodies, which might be less disruptive than uniform or random application. For example, snail control or behavioral modification programs might be directed at waterbodies that permit cattle use, because those waterbodies had significantly greater schistosome prevalence.

The higher prevalence of schistosome in waterbodies that permit cattle use, suggests quite effective community regulation of cattle-restricted ponds (*kisimas)* for maintaining access to safe water. Maintenance of these water sources is enforced by community members, and generally aims to keep waterbodies used for domestic and drinking water free from human and cattle excreta contamination. The specific rules and level of enforcement differ from community to community based on water availability and demand or knowledge of water borne diseases. Other studies have found that limiting waste disposal to water bodies serves as an alternative strategy for water borne disease control such as schistosomiasis (35,36). Thus, the targeted and consistent use of distinct waterbodies for distinct purposes could be a key behavioral intervention to reduce schistosome transmission until the broad availability of adequate water sanitation (37).The elevated schistosome prevalence in cattle permitted waterbodies could suggest that the schistosomes we observed in our study are *S. Bovis* or *S. haematobium or S. haematobium* x *S. bovis* hybrids directly using cattle or other livestock as definitive hosts (38). For example, a recent study in Benin identified cattle as potential definitive hosts for *S. haematobium* and *S. bovis*(39).

However, we would then also expect an elevated prevalence of nonschistosomes, which are likely predominantly infecting cattle,in cattle-permitted waterbodies.The equivalent prevalence of nonschistosome parasites regardless of cattle permission suggests that while humans may avoid contamination of *kisimas*, cattle regulations are not well adhered to. In practice, cattle in pastoral communities are not strictly managed, they are left wandering around even during flooding periods, which can lead to direct introduction or runoff into sites deemed as “restricted”(39). Spatially heterogeneously distributed infection of cattle and human schistosome parasites has been reported in other studies (40). Therefore, there is a growing need to pair eocological surveys, such as ours, with molecular identification techniques (41–43) to shed light on transmission dynamics of trematodes infecting humans, livestock, and wildlife within a One Health Approach. Resolving the genetic and species identities of the parasites we observed here could reveal direct interactions within snail vectors, e.g. dominance hierarchies (44), or clarify how coexisting parasite taxa differentially adapt to shared ecological gradients (45).While the prevalence of non-schistosomes did not increase with cattle permission, non-schistosome prevalence increased with waterbody size. This islikely due to the fact that larger waterbodies are used by more cattle, serve as cattle gathering stations and are less likely to dry up maximizing transmission in the later season.

Counter to our hypotheses, waterbody permanence and proximity to school were not significantly associated with snails or infections. We expected permanent waterbodies to be more suitable for snails and to facilitate human and animal access year-round, thereby having greater infection rates. Similarly, we expected that waterbodies nearby schools would be important exposure sites(46). The lack of association with school distance indicates that other activities may be more important, e.g., swimming, fishing, bathing, fetching water, searching water for watering livestock and washing clothes are among risk practices for schistosome transmission (47,48). Indeed, swimming during the weekend (i.e., non-schooldays) represents a substantial portion of water contact time for children(49). Similarly, a previous study found a positive spatial relationship between household schistosome infection rates and snail infection rates in the most proximate waterbodies, suggesting domestic, rather than school-based patterns of contact(20).Therefore, identification of specific transmission hotspots within a complex transmission network can serve as an alternative strategy for schistosomiasis control.

Snail presence and prevalence of nonschistosome parasites also varied between the two phases of the study, suggesting seasonal trends in trematode transmission(50). Specifically, moving from rainy to dry season, snail presence decreased, but nonschistosome prevalence increased. As the dry season progresses, snail populations decrease due to the contraction of waterbodies and the onset of snail aestivation (51). They also decrease due to increased disturbance caused by elevated human and livestock use (52,53). The increase in nonschistosome parasite prevalence could reflect the passage of adequate time for those parasites to complete their development and emerge from snail hosts. It could also reflect increased parasite input into these waterbodies as humans and livestock increasingly use fewer and fewer waterbodies as many become fully dry. Various other studies of snails and schistosomes demonstrate aligned and offset peaks in snail abundance and infection rates (17,40,50,54,55).Year-round studies with finer temporal resolution are needed in highly seasonal landscapes, such as this inland region of Northwestern Tanzania, to determine these seasonal patterns as well as how fluctuations in abiotic factors such as temperature, rainfall, water velocity influence snails and trematode transmission seasonality (56–58).

While we found little variation among villages or districts in snail presence or infection rates, we did not assess human, livestock, or wildlife infections. Therefore, our results do not imply that villages or districts do not vary substantially in the human or animal burden of infection or disease. Discrepancies in transmission between villages suggest that variation in water contact patterns is likely important. For example, household proximity to foci of snail vectors (20,59) and lack of local access to safe waters causes inevitable contact with infested water leading to high transmission(60).Therefore, our findings of waterbody-scale transmission factors can complement other studies across larger scales to illuminate hotspots within a network of complex transmission that can guide the development of targeted interventions for schistosomiasis control.

## Acknowledgments

The author acknowledges the support provided by Shinyanga, Mwanza and Simiyu region authorities and leadership of communities and villages involved in the survey. We thank Rashid Juma, BahatiLukomeso, Egbat Kazi, Revocatus Silayo, Francis Kagaruki, Revocatus Alphonce, James Kubeja and Amiru Swaibu for assistance in data collection. Dennis Byakuzana for driving the team to and from the field. We also acknowledge the National Research Ethical review committee of the National Institute for Medical research for approval of the study and National Institute of Health (NIH) for Funding.

## Author contributions

Conceived and designed research: TA NCS DJC SK. Performed research: TA NCS MPM JC DJC SK. Analyzed data: DJC NCS. Wrote the paper:TA NCS MPM JC DJC SK.

